# Opposing subclasses of *Drosophila* ellipsoid body neurons promote and suppress sleep

**DOI:** 10.1101/2021.10.19.464469

**Authors:** Abigail Aleman, Jaison Jiro Omoto, Prabhjit Singh, Bao-Chau Nguyen, Pratyush Kandimalla, Volker Hartenstein, Jeffrey M. Donlea

**Affiliations:** Department of Neurobiology, David Geffen School of Medicine at University of California – Los Angeles, Los Angeles, CA, 90095, United States; Molecular, Cellular & Integrative Physiology Interdepartmental Program, University of California – Los Angeles, Los Angeles, CA, 90095, United States; Department of Molecular, Cellular, and Developmental Biology, University of California – Los Angeles, Los Angeles, CA, 90095, United States

## Abstract

Recent work in *Drosophila* has uncovered several neighboring classes of sleep-regulatory neurons within the central complex. However, the logic of connectivity and network motifs remains limited by the incomplete examination of relevant cell types. Using a recent genetic-anatomical classification of ellipsoid body ring neurons, we conducted a thermogenetic screen to assess sleep/wake behavior and discovered two opposing populations: sleep-promoting R3m and wake-promoting R3d neurons. Activation of these neurons influences sleep duration and architecture by prolonging or shortening sleep bouts, suggesting a key role in sleep maintenance. R3m and R3d neurons are GABAergic and require GABA synthesis for their effects on sleep. Finally, we use a fluorescent reporter for putative synaptic partners to embed these neurons within the known sleep-regulatory network; R3m and R3d neurons lay downstream of wake-active Helicon cells, and R3m neurons likely inhibit R3d neurons. Together, the data presented herein suggest a neural mechanism by which previously uncharacterized circuit elements operate within the sleep homeostat to stabilize sleep-wake states.

## Introduction

Sleep is a state that is vital to maintain physiology and neural processing (Dongen et al., 2003; Everson, 1995; Everson et al., 1989; Shaw et al., 2002; Vaccaro et al., 2020) but the circuit organization of sleep control systems is not fully understood in any species. Previous studies in *Drosophila* have identified multiple neuronal populations in the central complex that contribute to a recurrent feedback loop to regulate sleep (Donlea et al., 2018; Liu et al., 2016). This circuit includes one subclass of ring neurons that innervate the anterior portion of the ellipsoid body (EB) (Liu et al., 2016), a toroid structure on the midline of the fly brain that also contributes to spatial orientation, navigation, motor control, and arousal (Bausenwein et al., 1994; Fisher et al., 2019; Kim et al., 2019; Kottler et al., 2019; Lebestky et al., 2009; Ofstad et al., 2011; Seelig and Jayaraman, 2015). The EB neuropil is comprised of several different cell populations, including strong innervation from several anatomically distinct subclasses of ring neurons (Hanesch et al., 1989; Lin et al., 2013; Omoto et al., 2018; Renn et al., 1999; Young and Armstrong, 2010), many of which can be subdivided into unique types based on morphology and connectivity criteria (Hulse et al., 2020). Most ring neurons are developmentally-related descendants of the DALv2/EBa1 lineage (R-neurons). Each R-neuron typically contains a dendritic arborization lateral to the EB, most commonly in the bulb, then projects a single axon in a concentric annulus within the EB neuropil (Hanesch et al., 1989; Omoto et al., 2018). The EB neuropil colocalizes with strong immunostaining for GABA (Enell et al., 2007; Hanesch et al., 1989; Shaw et al., 2018; Xie et al., 2017; Zhang et al., 2013) and is labeled by *gad1*-Gal4 (Enell et al., 2007; Kahsai and Winther, 2011), suggesting that many of the R-neurons that innervate EB domains may be inhibitory GABAergic neurons.

Recent experiments have identified an R-neuron subclass that elevates their electrical activity with waking experience and can drive a persistent increase in sleep upon thermogenetic activation (Liu et al., 2016). These neurons were initially referred to as R2 when their integrative role in sleep drive and placement within the sleep homeostat were described (Donlea et al., 2018; Liu et al., 2016), but have since been typically referred to as R5 (Blum et al., 2020; Omoto et al., 2018; Raccuglia et al., 2019) to avoid conflation with ring neurons that were historically designated R2 and play a role in feature detection (Hanesch et al., 1989; Omoto et al., 2017; Seelig and Jayaraman, 2013). Sleep- and arousal-related signals may reach EB R-neurons by at least two pathways: visual projection tubercular-bulbar (TuBu) neurons and arousal-encoding Helicon/ExR1 neurons (Donlea et al., 2018; Guo et al., 2018; Hulse et al., 2020; Lamaze et al., 2018; Raccuglia et al., 2019). Serotonergic modulation of other ring neuron populations can shape sleep architecture (Liu et al., 2019), but the precise roles of many R-neuron subclasses on sleep/wake regulation and organization have not been clearly described.

To test whether other R-neuron subclasses influence sleep regulation, we have used a thermogenetic activation screen using the warm-sensitive cation channel TrpA1 (Hamada et al., 2008). Our findings identify two R-neuron populations that influence sleep: sleep-promoting R3m cells and wake-promoting R3d neurons. Each of these populations colocalizes with GABA immunostaining, disruption of GABA synthesis in either population reduces sustained bouts of sleep or wake, and depletion of Gad1 in R3m neurons prevents flies from generating an acute increase in sleep following traumatic neural injury. To better understand the connectivity between R3m, R3d, and other sleep-regulatory cell types in the EB, we have imaged the distribution of putative contacts labelled by a genetically-encoded reporter for synaptic connections (Macpherson et al., 2015). We find that R3m and R3d neurons each likely lie downstream of arousal-promoting Helicon/ExR1 neurons and form few contacts with sleep-integratory R5 cells. Together, these results indicate that at least three R-neuron subclasses that project into the anterior portion of the EB (R3m, R3d, and R5) influence sleep regulation and that each population likely receives sleep/arousal cues via synaptic input from Helicon/ExR1 neurons.

## Results

### EB mini-screen for sleep-regulatory ring neurons

Previous studies have identified neuron types within the *Drosophila* central complex, including the R5 ring neurons of the ellipsoid body, that homeostatically promote sleep (Donlea et al., 2014; Liu et al., 2016; Pimentel et al., 2016). Whether additional circuitry also influences sleep-related signals in the EB, however, has not been clearly described. To examine the roles of other EB ring neuron subclasses in sleep regulation, we completed a targeted thermogenetic screen of genetic driver lines that drive expression in R-neuron subtypes (Jenett et al., 2012; Omoto et al., 2018; Tirian and Dickson, 2017). 27 genetic driver lines that label different subsets of R-neurons were each used to express the warm-sensitive cation channel TrpA1 (Hamada et al., 2008) **(Table 1)**. Experimental (Gal4/UAS-*TrpA1*) and genetic control flies (Gal4/+) for each driver were loaded into activity monitors for 1-2 days of baseline sleep, then heated to 31°C either for 6hr from ZT0-6 **(Figure 1A)** or for 12hr from ZT12-0 **(Figure 1B)**. Because sleep in wild-type flies is relatively low during the first few hours after ZT0 and high during the night, we hypothesized that the morning stimulation would enable us to preferentially identify sleep-promoting neurons while night-time activation would identify arousing R-neuron subclasses. R-neurons can be subdivided into at least 10 subclasses based on the locations of their dendritic arbors in the bulb and of their axonal projections into the EB **(See Schematic in Figure 1C)** (Hulse et al., 2020; Omoto et al., 2018). TrpA1 activation of many R-neuron subclasses, including R1, R2, R3a, R3p, and R4d, elicited little change in sleep time during either heat protocol. As described previously, activation of R5 neurons during either the day or night resulted in both acute and prolonged increases in sleep. Additionally, we also found that daytime stimulation of two genetic driver lines (*R28E01*-Gal4 and *R47D08*-Gal4 (Jenett et al., 2012)) labeling subsets of R3m neurons drove the largest acute increases in sleep **(Figures 1A, D)**. Conversely, two of the drivers that elicited the largest decrease in sleep during night-time stimulation, *R80C07*-Gal4 and *R84H09*-Gal4 (Jenett et al., 2012), localize their EB expression primarily to subpopulations of R3d ring neurons **(Figures 1B, D)**.

**Table 1.** Genetic drivers used to stimulate ring neuron subclasses, related to Figure 1. A list of the primary Ring neuron classes that are labelled by genetic drivers included in the TrpA1 activation mini-screens described in **Figure 1**.

| **Ring Neuron Class** | **Genetic Drivers** |
| --- | --- |
| R1 | *R31A12, VT016270* |
| R2 | *R78B06* |
| R3a | *R12G08* |
| R3d | *R80C07, R84H09, VT002226, VT019068, VT020613, VT029750* |
| R3m | *R28E01, R47D08, R28D01, R37E10, R92A09, VT0200036, VT042805* |
| R3p | *VT063740, VT063949* |
| R3w | *VT057232* |
| R4d | *R12B01* |
| R4m | *R59B10, R34D03, VT038873, VT026873* |
| R5 | *R58H05, VT004309* |

**Figure 1.**
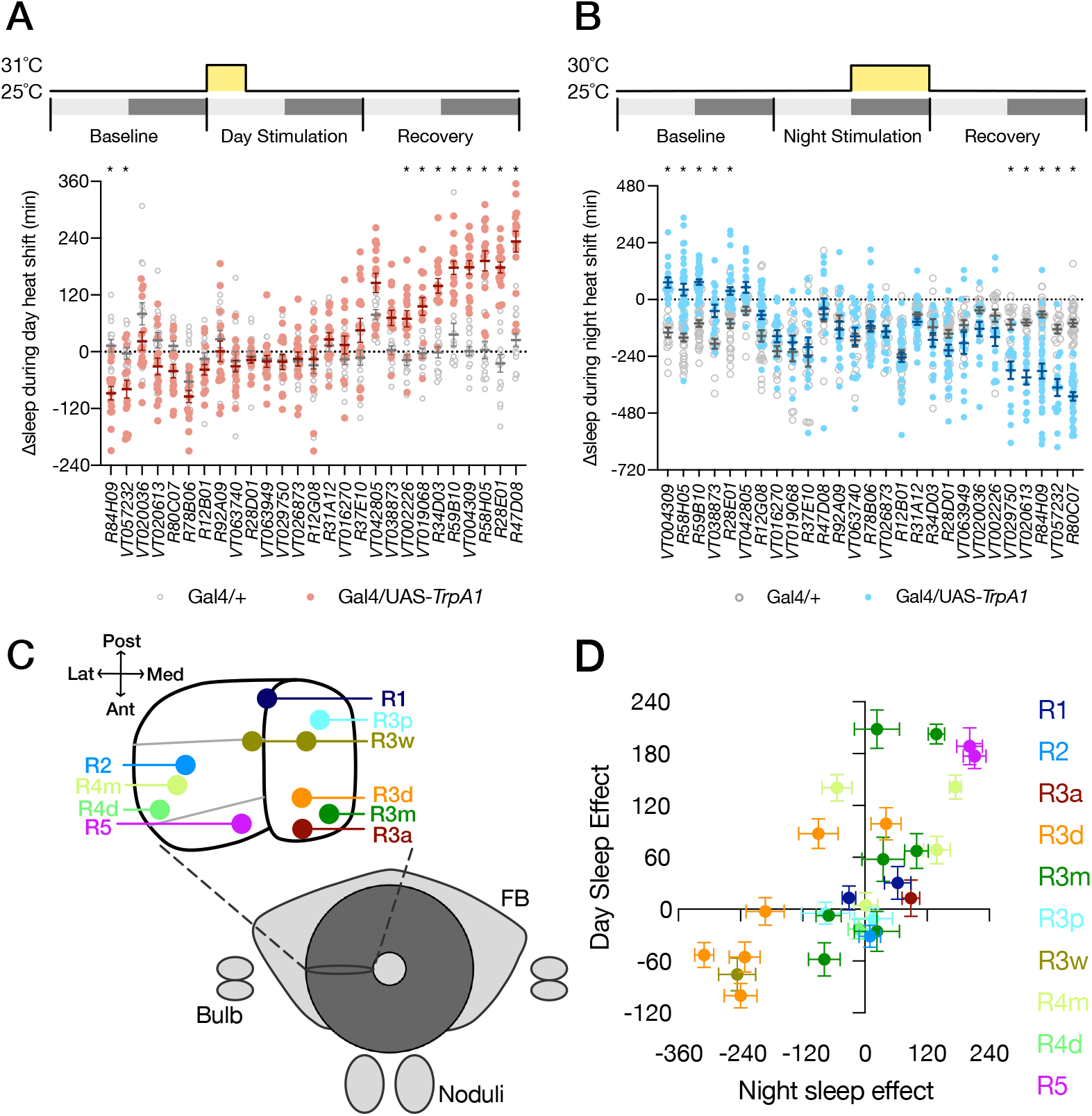
Characterizing sleep regulatory roles for distinct classes of Ring neurons. **(A-B)** Sleep results from targeted mini-screen for heat activation of Gal4 genetic drivers that label individual subtypes of EB Ring neurons. Change in sleep for each individual fly between heat exposure from ZT0-6 (**A**; 31°C) or ZT12-0 (**B**; 30°C) compared to sleep from the same fly during the previous baseline day. Gray depicts Gal4/+ controls for each driver; red or blue represents Gal4>UAS-*TrpA1* experimental lines. Significant Driver-by-Presence of *TrpA1* interactions were found for both day-time (F_(26,793)_=16.05, p<0.0001, n=14-32 flies/genotype) and night-time (F_(26,1110)_=18.49, p<0.0001, n=15-32 flies/genotype) mini-screens. * denotes drivers with p<0.05 by pairwise Šídák’s multiple comparisons test. **(C)** Schematic of the *Drosophila* Central Complex, including a frontal cross-section of the Ellipsoid body illustrating projection regions for 10 distinct Ring neuron subtypes. Based on (Omoto et al., 2018). **(D)** Scatter plot depicting the changes in sleep for each Gal4 driver included in the sleep mini-screen resulting from heat exposure during the night (x-axis) and day (y-axis). Day or night sleep effects were calculated by subtracting the Gal4/+ changes in sleep from the Gal4/TrpA1 changes in sleep shown in panels **A-B** for each driver. Datapoints are color coded by the Ring neuron subtypes in which each Gal4 driver is primarily expressed.

### Activation of R3m neurons acutely induces consolidated sleep

To precisely characterize the sleep-promoting effects of R3m neuron activation, we first confirmed the expression of *R28E01*-Gal4. Confocal projections included in **Figure 2A** indicate that *R28E01*-Gal4 expresses primarily in R3m neurons of the ellipsoid body, but also includes a small number of neurons in the suboesophageal zone and in the superior protocerebrum (Omoto et al., 2018). As shown in **Figure 2B**, heat stimulation acutely promotes sleep in *R28E01*-Gal4>UAS-*TrpA1* flies compared to genetic controls (*R28E01*-Gal4/+ and UAS-*TrpA1*/+). Over a 6h activation from ZT0-6, R3m activation using both *R28E01-*Gal4>UAS-*TrpA1* and *R47D08*-Gal4>UAS-*TrpA1* significantly increases both sleep time and mean sleep bout duration compared to genetic controls; no difference in either sleep time or bout length was detected between experimental flies and genetic controls on the baseline day prior to activation or recovery day following activation **(Figures 2C-D)**. R3m activation did not alter activity counts/waking minute **(Figure 2E)**, indicating that the gain in sleep that we observed was unlikely to be linked with changes in locomotor patterns. Because we observed an increase in the length but not the number of sleep bouts during R3m activation, we sought to test whether stimulating R3m neurons might preferentially enhance sleep maintenance with a weaker effect on the initiation of new sleep episodes. To test this possibility, we used a recently described behavioral analysis package to quantify the probability that a sleeping fly would awaken (P(wake)) or that a waking fly would fall asleep in each minute (P(doze)) (Wiggin et al., 2020). As shown in **Figures 2F-G**, R3m stimulation using *R28E01*-Gal4 or *R47D08*-Gal4 (dark green), resulted in a decreased P(wake) compared both to genetic controls that were also housed at 31°C or to data from the experimental flies that was collected either on the baseline day or recovery day. Experimental *R28E01-*Gal4/UAS-*TrpA1* flies showed little change in P(doze) across the three experimental days while *R47D08*-Gal4*/*UAS-*TrpA1* flies had a significant increase in P(doze) during heat exposure **(Figures 2F, H)**. These data indicate that R3m activation has mixed effects on the initiation of sleep bouts, but strongly suppresses the minute-by-minute probability that a sleeping fly will wake up.

**Figure 2.**
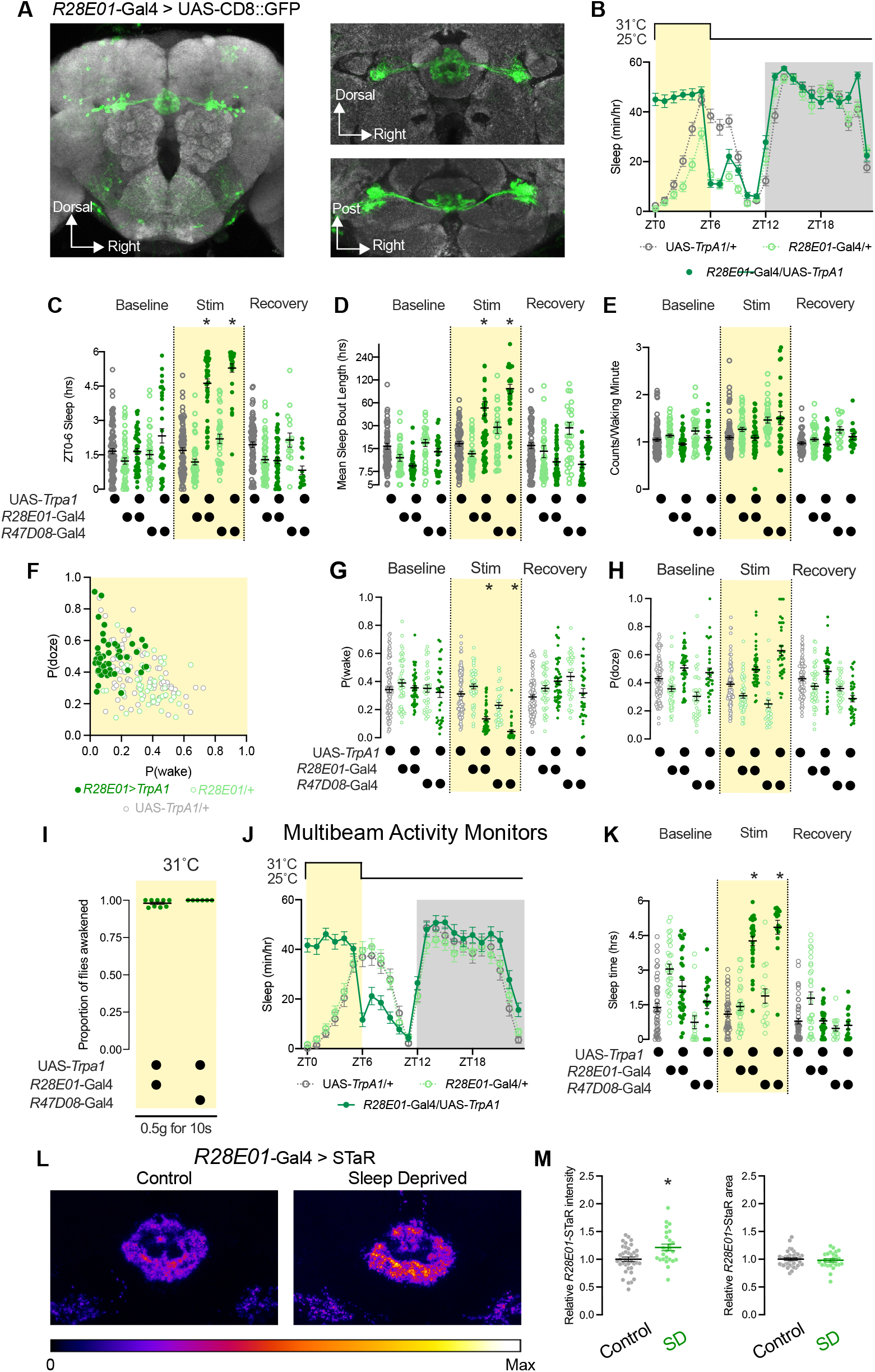
R3m neuron activity acutely increases sleep time and consolidation. **(A)** Expression of CD8::GFP under the control of *R28E01*-Gal4 localizes to the R3m subpopulation of EB Ring neurons. Left, z-projection of whole brain confocal scan; Right, Central complex expression of *R28E01*-Gal4 as projections of higher magnification confocal scans from frontal (top) or dorsal (bottom) views. **(B)** Hourly sleep timecourse of *R28E01-*Gal4>UAS-*TrpA1* flies (dark green) compared to *R28E01-*Gal4/+ (light green) and UAS-*TrpA1*/+ controls (gray) during the day of a 6-h heat stimulation (31°C) from ZT0-6. Two-way repeated measures ANOVA finds a significant Time-by-Genotype interaction (F_(46, 3496)_ = 29.46, p<0.0001, n=48-59 flies/group). **(C-D)** Sleep time **(C)** and average sleep bout length **(D)** from ZT0-6 is increased in *R28E01-* Gal4>UAS-*TrpA1* and *R47D08-*Gal4>UAS-*TrpA1* flies (dark green) compared to genetic controls (*R28E01-*Gal4/+ and *R37D08*-Gal4/+ shown in light green; UAS-*TrpA1*/+ in gray) during heat stimulation. No significant difference between experimental (dark green) and genetic controls (light green or gray) could be detected on either baseline or recovery days. Two-way repeated measures ANOVA finds a significant Day-by-Genotype interaction for both sleep time (F_(8, 478)_ = 78.15, p<0.0001, n=30-88 flies/group) and bout length (F_(8, 478)_ = 21.99, p<0.0001, n=30-88 flies/group). **(E)** Mean counts/waking minute from ZT0-6 is not changed in *R28E01-*Gal4>UAS-*TrpA1* and *R47D08-*Gal4>UAS-*TrpA1* flies (dark green) during heat stimulation compared to genetic or temperature controls. **(F)** Scatter plot depicting the probability that a sleeping fly will awaken (P(wake); X-axis) and the probability that a fly will transition from wake to sleep (P(doze); Y-axis) for individual animals during a 6-h heat exposure. Dark green shows experimental *R28E01-*Gal4/UAS-*TrpA1* flies; *R28E01-*Gal4/+ controls in light green and UAS-*TrpA1*/+ in gray. **(G-H)** P(wake) **(G)** and P(doze) **(H)** values during ZT0-6 for baseline, heat stimulation, and recovery days for experimental flies (*R28E01*-Gal4/UAS-*TrpA1* and *R47D08*-Gal4/UAS-*TrpA1*; dark green) and genetic controls (UAS-*TrpA1*/+ in gray; *R28E01*-Gal4/+ and *R47D08*-Gal4/+ in light green). Two-way repeated measures ANOVA finds a significant Day-by-Genotype effect for P(wake) (F_(8, 478)_=23.90, p<0.0001, n=30-88 flies/group) and P(doze) (F_(8, 478)_=23.77, p<0.0001, n=30-88 flies/group). **(I)** Mechanical vibration (0.5g for 10s) was sufficient to wake both *R28E01*-Ga4/UAS-*TrpA1* and *R47D08*-Gal4/UAS-*TrpA1* flies from sleep when housed at 31°C. **(J)** Multibeam monitors detect increased sleep in *R28E01-*Gal4>UAS-TrpA1 flies (dark green) during heat exposure (31°C) from ZT0-6 compared to *R28E01-*Gal4/+ (light green) and UAS-*TrpA1*/+ (gray) genetic controls. Two-way repeated measures ANOVA finds a significant Time-by-Genotype interaction (F_(46, 2070)_ = 16.58, p<0.0001, n=30-32 flies/group). **(K)** Total sleep time from ZT0-6 is increased in both *R28E01*-Gal4>UAS-*TrpA1* and *R47D08*-Gal4>UAS-*TrpA1* flies (dark green) compared to Gal4/+ (light green) and UAS-*TrpA1*/+ (gray) controls during heat exposure (yellow shading), but not during baseline or recovery days. **(L)** Example images from sleep deprived (right) and control brains (left) showing BRP::smFP_V5 labelling of pre-synaptic active zones in R3m neurons using *R28E01*-Gal4. **(M)** Quantification of relative STaR intensity (left panel) and cross-sectional area (right panel) for groups shown in 2I. Sleep deprived brains showed an increase in R3m>STaR intensity (Two-tailed T-test p=0.0031; n=25-37 each group), but no change in area (Two-tailed T-test p=0.5804; n=25-37 each group).

To test whether the increased quiescence observed during R3m activation fits the same behavioral criteria as spontaneous sleep, we next tested arousability. Nearly all experimental flies were awakened by a brief vibrational stimulus delivered to their activity monitors (0.5g for 10s), consistent with the idea that R3m activation drives a rapidly reversible sleep state **(Figure 2I)**. Next, we tested the effects of R3m activation using multi-beam activity monitors, which provide more precise measurements of movement and positional data than single-beam monitors. Experimental flies exhibited similar increases in quiescence using both systems **(Figures 2J-K)**, confirming that the sleep induction that we observed in single-beam monitors was not confounded by micromovements or other non-sleep behaviors. In contrast to the prolonged sleep increase that has been previously described following R5 activation (Liu et al., 2016), R3m stimulation elicited only an acute increase in sleep during heat exposure which rapidly dissipated when flies were returned to 25°C **(Figure 2B)**. To test whether sleep-promoting R3m neurons might exhibit structural plasticity in response to sleep loss, we expressed a flp-based reporter for the active zone protein Bruchpilot (BRP), Synaptic Tagging with Recombination (STaR), using *R28E01*-Gal4 (Chen et al., 2014; Kittel et al., 2006; Peng et al., 2018). As shown in **Figures 2L-M**, R3m neurons from sleep deprived flies increased their pre-synaptic BRP::smFP_V5 by 21.2 ±6.875% compared to those in rested siblings. R3m neurons, therefore, may increase their synaptic release to trigger sleep at times of heightened sleep need.

### R3d neuron stimulation promotes physiological wakefulness

Our ring neuron mini-screen shown in **Figure 1** indicated that two Gal4 drivers that primarily label R3d neurons, *R80C07-*Gal4 **(See Figure 3A)** and *R84H09*-Gal4, promote wakefulness upon overnight activation with *TrpA1*. The reduction in sleep that occurs during R3d activation **(Figures 3B-C)** coincides with a fragmentation in sleep bout length **(Figure 3D)**, but no change in bout number **(Figure 3E)**. R3d stimulation using *R80C07-*Gal4 also caused a modest decrease in activity counts per waking minute **(Figure S1A)**, suggesting that R3d neurons do not drive hyperarousal. Sleep loss during R3d activation with *R80C07*-Gal4 or *R84H09*-Gal4 can be attributed to a reduction in the persistence of sleep bouts; the probability that a sleeping fly will awaken, P(wake), is significantly elevated with R3d activation compared to genetic controls **(Figures 3F-G)**, while the likelihood of sleep initiation, P(doze), is unaffected by R3d stimulation **(Figures 3F, H)**. Unlike in R5 and R3m neurons, STaR labelling shows no significant change in the active zone abundance of R3d neurons after overnight sleep loss **(Figure S1B)**, consistent with previous results indicating that pre-synaptic upscaling may not occur across all R-neuron subclasses during sleep deprivation (Liu et al., 2016).

**Figure 3.**
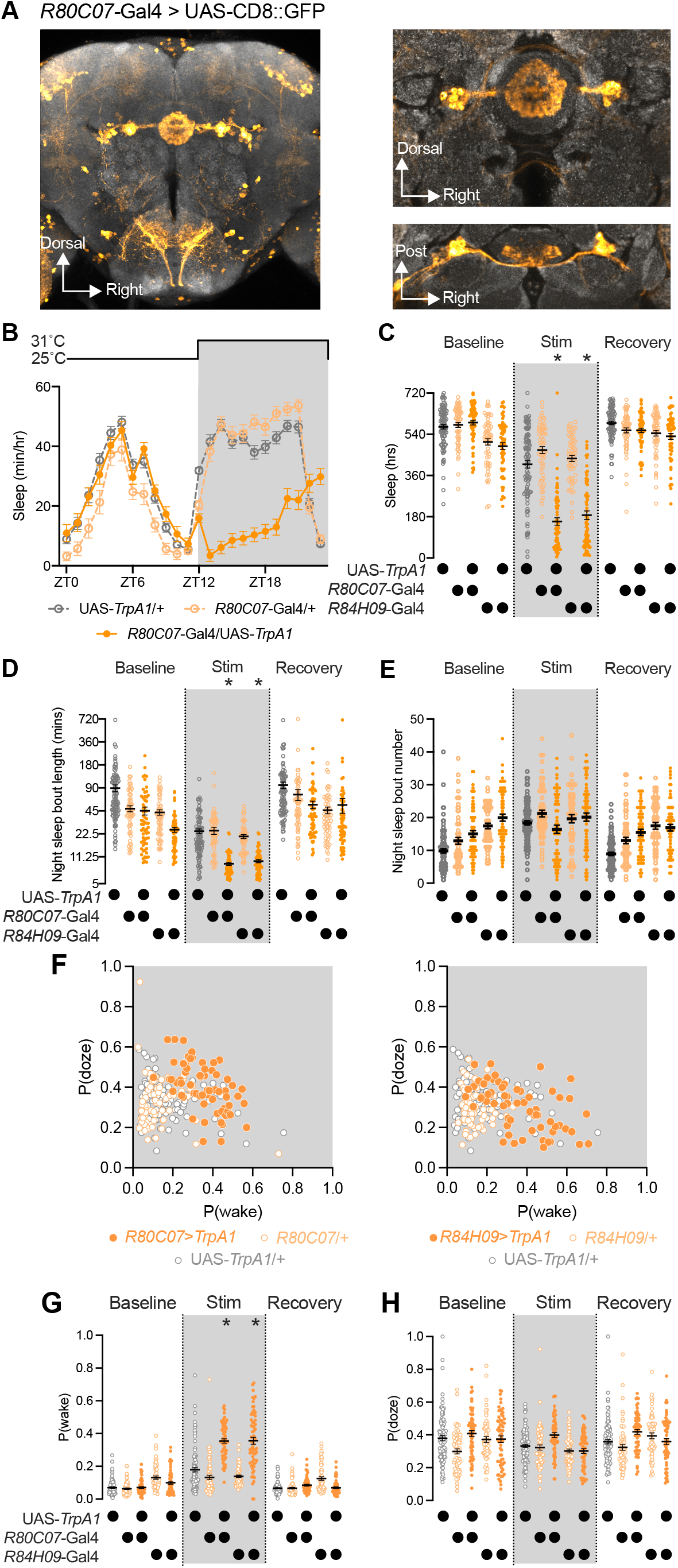
R3d neuron activation promotes waking via reduced sleep maintenance. **(A)** Expression of CD8::GFP under the control of *R80C07-*Gal4 is primarily localized to R3d ring neurons. Left, z-projection of whole brain confocal scan; Right, Central complex expression of *R80C07*-Gal4 shown as projections of higher magnification confocal scans from frontal (top) or dorsal (bottom) angles. **(B)** Hourly sleep time-course of *R80C07*-Gal4/UAS-*TrpA1* experimental flies (dark orange) compared to *R80C07*-Gal4/+ (light orange) and UAS-*TrpA1*/+ (gray) controls during a 12-h overnight heat exposure (30°C; yellow shading) from ZT12-24. Two-way repeated measures ANOVA finds a significant Time-by-Genotype interaction (F_(46, 2875)_ = 33.26, p<0.0001, n=32-64 flies/group). **(C-D)** Night-time sleep **(C)** and average sleep bout duration **(D)** are decreased in *R80C07*-Gal4>UAS-*TrpA1* and *R84H09*-Gal4>UAS-*TrpA1* flies (dark orange) compared to Gal4/+ (light orange) and UAS-*TrpA1*/+ (gray) controls. Two-way repeated measures ANOVA finds a significant Day-by-Genotype interaction for sleep time (F_(8, 668)_ = 71.81, p<0.0001, n=59-93 flies/group) and for average night-time sleep bout length (F_(8, 668)_ = 4.593, p<0.0001, n=59-93 flies/group). **(E)** R3d stimulation in *R80C07*-Gal4>UAS-*TrpA1* or *R84H09*-Gal4>UAS-*TrpA1* flies (dark orange) does not significantly alter night-time sleep bout number compared to genetic controls (light orange or gray). Two-way repeated measures ANOVA finds a significant Day-by-Genotype interaction (F_(8, 668)_ = 15.48, p<0.0001, n=59-93 flies/group). **(F)** Scatter plots showing individual fly values for P(wake) and P(doze) values during overnight (ZT12-0) heat exposure to stimulate R3d neurons; *R80C07-*Gal4 activation shown in left panel and *R84H09*-Gal4 used on right. The same UAS-*TrpA1*/+ flies (gray) are shown in both graphs. **(G-H)** P(wake) (G) and P(doze) (H) for R3d activation; experimental flies (dark orange) show increased P(wake) during overnight heat exposure compared to genetic controls (Gal4/+ groups in light orange and UAS-*TrpA1*/+ in gray), while no decrease in P(doze) was detected. Two-way repeated measures ANOVA finds a significant Day-by-Genotype interaction for P(wake) (F_(8,665)_ = 75.76, p<0.0001, n=58-93) and P(doze) (F_(8,665)_ = 4.686, p<0.0001, n=58-93).

### R3m and R3d influence sleep via GABA signaling

Previous reports found strong immunolabeling for GABA in the ellipsoid body, suggesting that many ring neurons are likely GABAergic (Hanesch et al., 1989; Xie et al., 2017; Zhang et al., 2013). As depicted in **Figures 4A-B**, our histology confirmed overlap between anti-GABA immunostaining and most, but not all, *R28E01*- and *R80C07-*positive cell bodies. We tested the effect of disrupting GABA production on sleep/wake regulation by expressing two independent RNAi constructs targeting the GABA synthesis enzyme Gad1 in R3m and R3d neurons (Dietzl et al., 2007; Jackson et al., 1990). While RNAi knock-down of Gad1 in R3m neurons using *R28E01*-Gal4 with both effector constructs reduced sleep during the night, R3d expression of either Gad1 RNAi had little effect on sleep **(Figure 4D)**. Further, suppressing *gad1* expression in R3m neurons altered sleep architecture by reducing the mean length of night-time sleep bouts **(Figure 4E)**. Expression of either *gad1* RNAi construct in R3m neurons also increased night-time P(wake) **(Figure S2)**, supporting the hypothesis that R3m electrical activity and GABA release promotes the persistence of consolidated sleep bouts. Conversely, expression of Gad1 RNAi constructs in R3d neurons did not alter sleep consolidation during the day or the night, but one of the two Gad1 RNAis shortened the length of daytime wake bouts when driven with *R80C07*-Gal4 **(Figure 4F)**. It is unclear whether the differing effects of Gad1 RNAi in R3ds may be attributable to variable efficacy of the constructs or to other factors, but the results support a possible role for GABA release from R3d neurons in stabilizing prolonged episodes of wakefulness. Next, we investigated whether reducing GABA signaling in sleep-promoting R3m neurons could prevent flies from elevating sleep time under conditions of high sleep pressure by testing responses to antennal injury (Singh and Donlea, 2020). In *R28E01-*Gal4/+ and UAS-*Gad1*^RNAi^/+ controls, flies showed large increases in sleep during the several hours after antennectomy **(Figures 4G-H, gray and light green)**. Experimental flies expressing RNAi constructs for *gad1* using *R28E01*-Gal4, in contrast, showed no significant increase in sleep during the day of antennal injury **(Figures 4G-H, dark green)**, indicating that GABA signaling from R3m is required for sustained sleep bouts on a nightly basis and to acutely promote sleep in response to traumatic axotomy.

**Figure 4.**
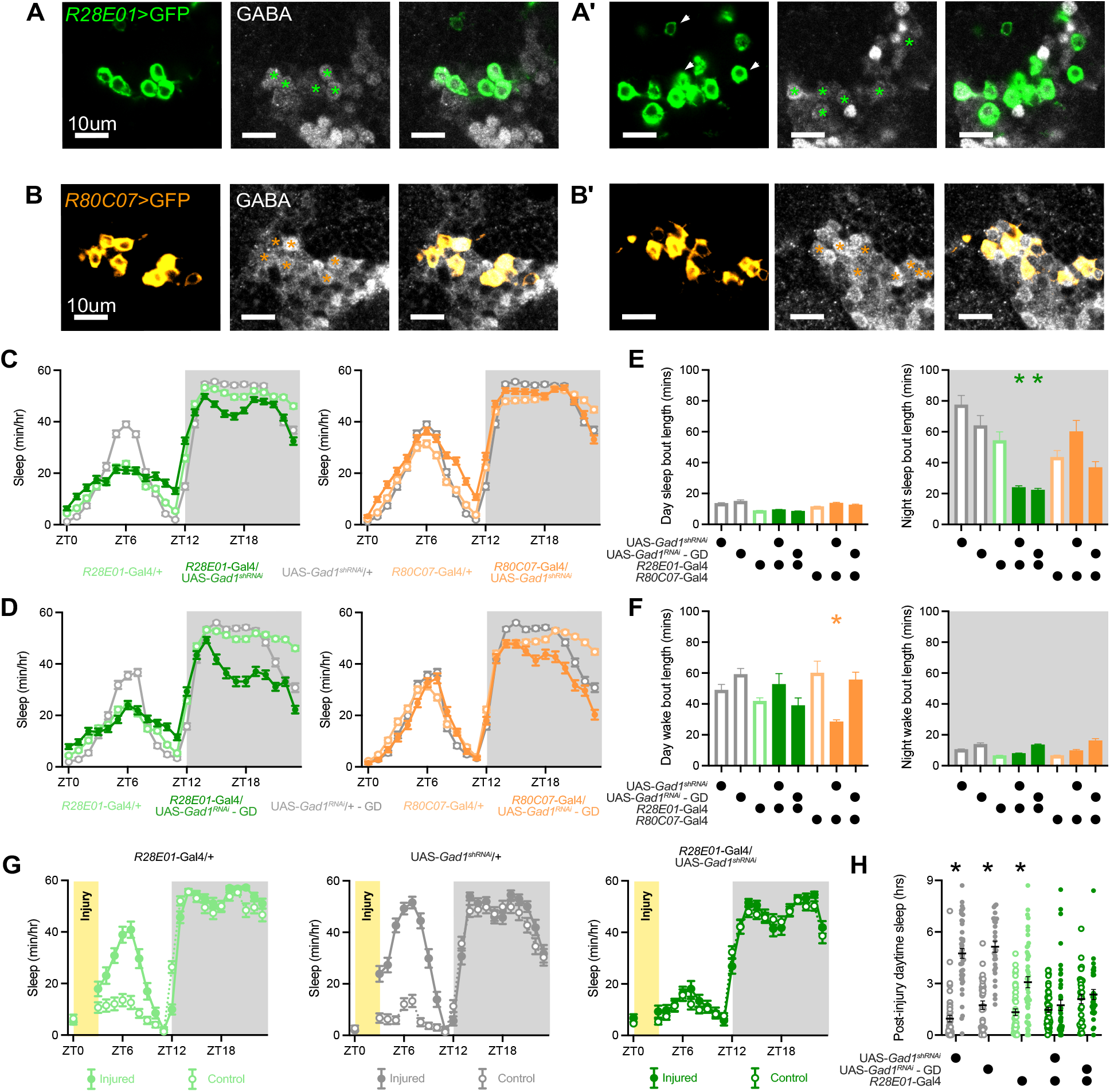
GABA synthesis in R3m and R3d neurons influences sleep regulation. **(A)** *R28E01*-positive R3m soma colocalize with immunostaining for GABA. Panels A and A’ represent distinct confocal slices within an individual z-stack. Asterisks indicate GABA-positive R3m neurons, while arrows represent R3m cell bodies with no detectable GABA signal. **(B)** R3d neurons labelled with *R80C07*-Gal4>UAS-CD8::GFP overlap with immunostaining for GABA. Panels B & B’ depict different confocal slices from an individual z-stack. Asterisks mark Gal4-labeled R3d soma which express GABA. **(C-D)** Hourly sleep timecourse for RNAi-mediated knock-down of Gad1 in R3m neurons using *R28E01*-Gal4 (left panels; green) and in R3d neurons expressing *R80C07-*Gal4 (right panels; orange). Sleep while expressing UAS-*Gad1*^shRNA^ SH330039 shown in **(C)**; UAS-*Gad1*^RNAi^ stock GD32344 shown in **(D)**. **(E)** Quantification of mean sleep bout length during the day (left panel) and night (right panel) for groups shown in **(C-D)**. One-way ANOVA finds a significant effect of genotype during the day (F_(7, 1221)_=17.01, p<0.0001, n=92-191) and night (F_(7, 1222)_=11.68, p<0.0001, n=92-192 flies/group). **(F)** Quantification of mean wake bout length during the day (left panel) and night (right panel) for groups shown in **(C-D)**. One-way ANOVA finds a significant effect of genotype during the day (F_(7, 1221)_=4.459, p<0.0001, n=92-191) and night (F_(7, 1222)_=20.94, p<0.0001, n=92-192 flies/group). **(G)** Hourly sleep time-course for experimental *R28E01-*Gal4>UAS-*Gad1*^shRNA^ (right panel; dark green) and control *R28E01*-Gal4/+ (left panel; light green) and UAS-*Gad1*^shRNA^/+ (middle panel; gray) on the day of antennectomy. Yellow shading depicts timing of antennal injuries. **(H)** Daytime sleep during the hours immediately following antennectomy (ZT4-12) for the groups shown in **(G)**. Two-way ANOVA finds a significant genotype-by-injury interaction (F_(4,235)_=16.12, p<0.0001, n=14-32 flies/group).

### R3m and R3d sleep-regulatory neurons lay downstream of Helicon/ExR1 neurons

To test whether R3m and R3d neurons might interact with other sleep-regulatory circuitry in the EB, we imaged the proximity of these cells to Helicon/ExR1 and R5 neurons, which both regulate sleep (Donlea et al., 2018; Hanesch et al., 1989; Liu et al., 2016; Omoto et al., 2018). Neurites from each of these four cell populations (R3m, R3d, Helicon/ExR1, and R5) closely neighbor each other, with all innervating the anterior portion of the EB and domains of the bulb **(Figure 5)**. Next, we used a genetic reporter for synaptic contacts, GFP Reconstitution Across Synaptic Partners (GRASP) (Feinberg et al., 2008; Macpherson et al., 2015), to test whether each pair of neurons forms putative synaptic contacts. For these studies, we used an activity-dependent GRASP variant that fuses one portion of GFP to the pre-synaptic vesicle protein nSyb (DiAntonio et al., 1993)while the remaining GFP epitopes were targeted to the plasma membrane of candidate post-synaptic partners, thereby labeling recently active points of contact across the synaptic cleft (Macpherson et al., 2015). Despite the close physical proximity of R3m, R3d, R5, and Helicon/ExR1, robust GRASP signal in the EB could only be found for Helicon/ExR1 → R3m and Helicon/ExR1 → R3d connections, while weaker signal was detected for R3m → R3d contacts **(Figures 5A, C, E)**. We also expressed a genetically encoded anterograde tracer, trans-Tango (Talay et al., 2017), in Helicon/ExR1 neurons and found that labelling was restricted solely to ring neurons of the anterior and inner central domains of the EB, which is consistent with direct connections from Helicon/ExR1 to R5, R3m, and R3d **(Supplemental Figure S3)** (Omoto et al., 2018). GRASP labelling for R3m → R3d and, more weakly, for R3m → Helicon/ExR1 contacts were also detected in the bulb **(Figures 5A, E)**, consistent with previous trans-Tango tracing for R3m targets (Omoto et al., 2018). No reliable signal was found to report contacts between R5 and either R3m or R3d **(Figures 5B, D)**. These data closely parallel connectivity patterns revealed by detailed reconstructions from serial electron micrographs (Hulse et al., 2020; Scheffer et al., 2020), patch clamp electrophysiology (Donlea et al., 2018), and genetic tracing experiments (Omoto et al., 2018). As schematized in **Figure 5F**, these results indicate that Helicon/ExR1 neurons relay sleep and arousal related cues directly to sleep-promoting R5 and R3m cells along with sleep-inhibiting R3d neurons to influence sleep timing and architecture, and that R3m neurons may suppress wake-promoting Helicon/ExR1 and R3d cells to maintain sleep bouts.

**Figure 5.**
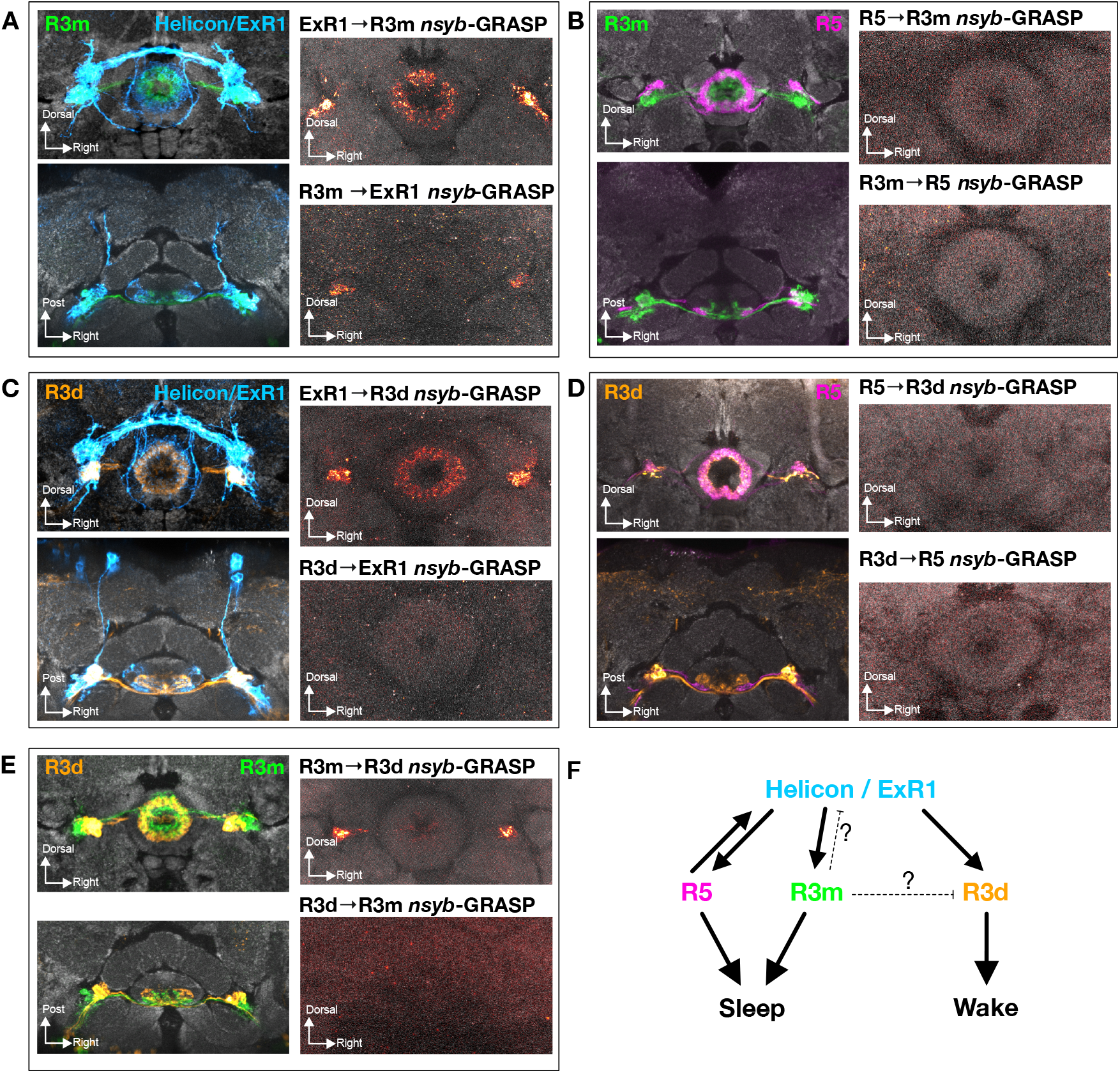
Connectivity between sleep-regulatory neuron types of the Ellipsoid body. **(A-B)** R3m neuron processes (green) neighbor neurites from both Helicon/ExR1 (A, left panels; blue) and R5 (B, left panels; magenta). *nsyb-*GRASP expression reports putative synaptic contacts from Helicon/ExR1 to sleep-promoting R3m neurons (A, top right panel), but not from R5 to R3m (B, top right panel). *nsyb*-GRASP fluorescence suggests R3m to Helicon/ExR1 contacts in the bulb (A, bottom right panel). Little, if any, signal was found to indicate synaptic connections from R3m to R5 (B, bottom right panel). **(C-D)** Left panels show frontal (top) and dorsal (bottom) views of R3d (orange) neurite localization in the proximity of Helicon/ExR1 (C, blue) and R5 (D, magenta) neurons. Neurites from both cell types are in close proximity in the bulb and EB and *nsyb-*GRASP labelling shows putative synaptic contacts from ExR1 to R3d neurons (C, top right panel), but we found no GRASP signal from R5 to R3d cells (D, top right). No evidence was found for connections from R3d to Helicon/ExR1 (C, bottom right) or R5 (D, bottom right) *nsyb-*GRASP. **(E)** Left; R3m (green) and R3d (orange) innervate closely neighboring zones of the EB and bulb. Using *nsyb*-GRASP, we found evidence for R3m to R3d synaptic contacts (top right panel), but no signal representing R3d to R3m connections (bottom right panel). **(F)** Schematic of putative synaptic connectivity between sleep-regulatory Helicon/ExR1, R5, R3m, and R3d neurons. This cartoon is based on our GRASP results (A-E) along with previously described electrophysiological studies (Donlea et al., 2018) and circuit reconstruction from serial electron micrographs (Hulse et al., 2020; Scheffer et al., 2020).

## Discussion

Our TrpA1 activation screen identifies novel roles for two R-neuron subtypes, R3m and R3d, in sleep regulation. Thermogenetic activation of R3m neurons acutely increased sleep via sustaining consolidated sleep bouts and suppressing the likelihood that flies would awaken. Conversely, R3d activation suppresses sleep by impairing sleep maintenance and increasing the probability of awakening without resulting in locomotor hyperactivity. Both R3m and R3d receive putative synaptic contacts from arousal-encoding Helicon/ExR1 neurons, but nSyb-GRASP reports few, if any, contacts from sleep-integratory R5 cells to R3m or R3d neurons under rested conditions. Both R3m and R3d colocalize with immunostaining for GABA, consistent with previous reports of strong GABAergic staining across the EB (Hanesch et al., 1989; Shaw et al., 2018; Xie et al., 2017; Zhang et al., 2013), and disrupting GABA synthesis in R3m neurons reduces total night-time sleep, impairs sleep consolidation at night, and prevents flies from increasing their sleep in response to neural injury.

Although the axons of R5, R3m, and R3d neurons are closely adjacent, our GRASP studies suggest that they are unlikely to be tightly interconnected within the EB. We found no evidence for connections between R5 and either R3m or R3d, or from R3d to R3m neurons. While GRASP reported possible contacts from R3m to R3d in the bulb, our STaR labelling of presynaptic BRP in R3m neurons showed little indication of active zones in the bulb. These results are complementary with findings from recent synaptic tracing and connectomics studies (Hulse et al., 2020; Omoto et al., 2018; Scheffer et al., 2020) and suggest that direct synaptic connections between classes of sleep-regulatory ring neurons may be limited. R5, R3m, and R3d each receive synaptic connections from Helicon/ExR1 neurons but, aside from possible R3m to R3d contacts, appear to form few contacts across ring neuron types (Donlea et al., 2018; Hulse et al., 2020; Omoto et al., 2018). Intriguingly, both R5 and R3m neurons display structural plasticity after prolonged wakefulness (Liu et al., 2016); while these neurons may be strengthening their outputs to existing synaptic partners, it is alternatively possible that transient synapses may be forming with novel partners in a context-dependent manner. Daily plasticity in the circadian clock, for instance, changes the neuron types that form synaptic connections with core pacemaker neurons from the morning to the evening (Fernandez et al., 2020; Song et al., 2021), indicating that circuit reconfiguration across circadian time or waking experience can influence sleep/wake regulation. Future investigations may probe the potential reconfiguration of ring neuron connectivity depending on sleep history. Furthermore, it remains unclear whether these ring neuron subtypes communicate via non-synaptic means, such as volume transmission or gap junctions, act in series via an intermediary, or form parallel sleep regulatory modules.

Interestingly, while activation of R3m and R5 neurons each increases sleep, the somnogenic effects of R3m activation rapidly dissipate once animals return to baseline while R5 activation results in a longer-lasting increase in sleep pressure (Liu et al., 2016). These differences suggest that R3m and R5 are unlikely to be functionally redundant but may instead comprise distinct components of a sleep control system or drive separate sleep stages, like the neighboring but separable circuits that influence REM and non-REM sleep in mammals (Anaclet et al., 2014; Chowdhury et al., 2019; Hayashi et al., 2015; Weber et al., 2015; Yu et al., 2019). Evidence for distinct neurophysiological activity patterns (Tainton-Heap et al., 2021; Yap et al., 2017) and varied arousal thresholds (Alphen et al., 2013) across *Drosophila* sleep bouts supports the possibility that *Drosophila* exhibit distinct sleep stages that might be regulated by different circuitry. Future studies will examine whether R3m and R5 promote sleep via parallel pathways, possibly via common downstream effector circuitry, and whether their activity is mutually exclusive with that of wake-promoting R3d cells. Newly developed preparations to image neural activity over many hours or days may open future investigations into how each of these neuron classes contributes daily sleep-wake patterns (Huang et al., 2018; Liang et al., 2016, 2019; Tainton-Heap et al., 2021; Valle et al., 2021).

Analysis of sleep/wake transition probabilities indicate that R3m and R3d activation influence sleep time by having opposing effects on sleep maintenance; activation of R3m neurons drives long periods of consolidated sleep, while R3d stimulation results in fragmented sleep episodes. In these roles, R3m and R3d each act to promote or oppose sleep maintenance with little effect on the initiation of sleep episodes. Because R3m neurons show increased BRP abundance after sleep loss and impairing GABA synthesis in R3m neurons prevents flies from increasing their sleep following neural injury, it is likely that R3m cells may engage at times of high sleep need to promote consolidated sleep. Further investigations will be required to identify the precise conditions in which R3d neurons promote waking, but RNAi-mediated knock-down of Gad1 in R3d neurons using a short-hairpin construct impairs the maintenance of wake episodes during the daytime. While serotonergic and dopaminergic release onto a collection of ring neuron types can shape sleep architecture and arousal (Lebestky et al., 2009; Liu et al., 2019), our data indicate that acute activation of specific ring neuron types can drive potent sleep changes. The sleep-regulatory ring neurons that we examine here may be included in the serotonin- and/or dopamine-responsive cell types that are known to influence sleep consolidation or arousability, but the precise physiological effects of neuromodulators on each cell type remain to be examined. The opposing sleep-regulatory effects of R3m and R3d neurons, in combination with R5 neurons, suggest that parallel representations of sleep- and wake-drive may be maintained between different populations of neurons in the EB. These three ring neuron types may represent the primary synaptic targets of arousal-encoding Helicon/ExR1 neurons (Hulse et al., 2020; Omoto et al., 2018), and control of their activity provides new opportunities to probe the organization of sleep control circuitry in the fly. While the downstream targets of R3m, R3d, and R5 neurons remain to be characterized, these cells lay in a promising intersection to relay arousal state signals from sleep control neurons to circuits that encode spatial representations of the exterior world (Donlea et al., 2018; Green et al., 2017; Kim et al., 2017, 2019; Liu et al., 2016; Seelig and Jayaraman, 2013, 2015).

## Acknowledgements

We thank all members of the Donlea lab for helpful discussions and feedback during this project, especially Jackie Weiss for technical assistance with histology protocols. Fly stocks were generously provided by Drs. Orkun Akin (UCLA), Gero Miesenbock (University of Oxford), and Paul Shaw (Washington University in St. Louis), Bloomington Drosophila Stock Center, the HHMI Janelia Research Campus, and the Vienna *Drosophila* Research Center. This project was supported by an Early Career Development Award from the Sleep Research Society Foundation to JD, a Career Development Award from the Human Frontiers Science Program to JD (CDA00026-2017-C), a Neuroscience Fellowship from the Klingenstein and Simons Foundations, NIH grant NS105967 to JD, an A.P. Giannini Postdoctoral Fellowship to JO, and a Cota-Robles Scholarship from UCLA to AA.

## Author Contributions

Conceptualization: J.O. & J.D., Methodology: J.O, A.A., P.S., Investigation: J.O., A.A., P.S., B-C. N, P.K., J.D., Writing – Original Draft: J.D., A.A., P.S., Writing – Review & Editing: J.O., A.A., P.S., B-C. N, P.K., V.H., J.D., Funding Acquisition – J.D., J.O. & V.H., Supervision – J.D.

## Declaration of Interests

The authors declare no competing interests

**Supplemental Figure 1.**
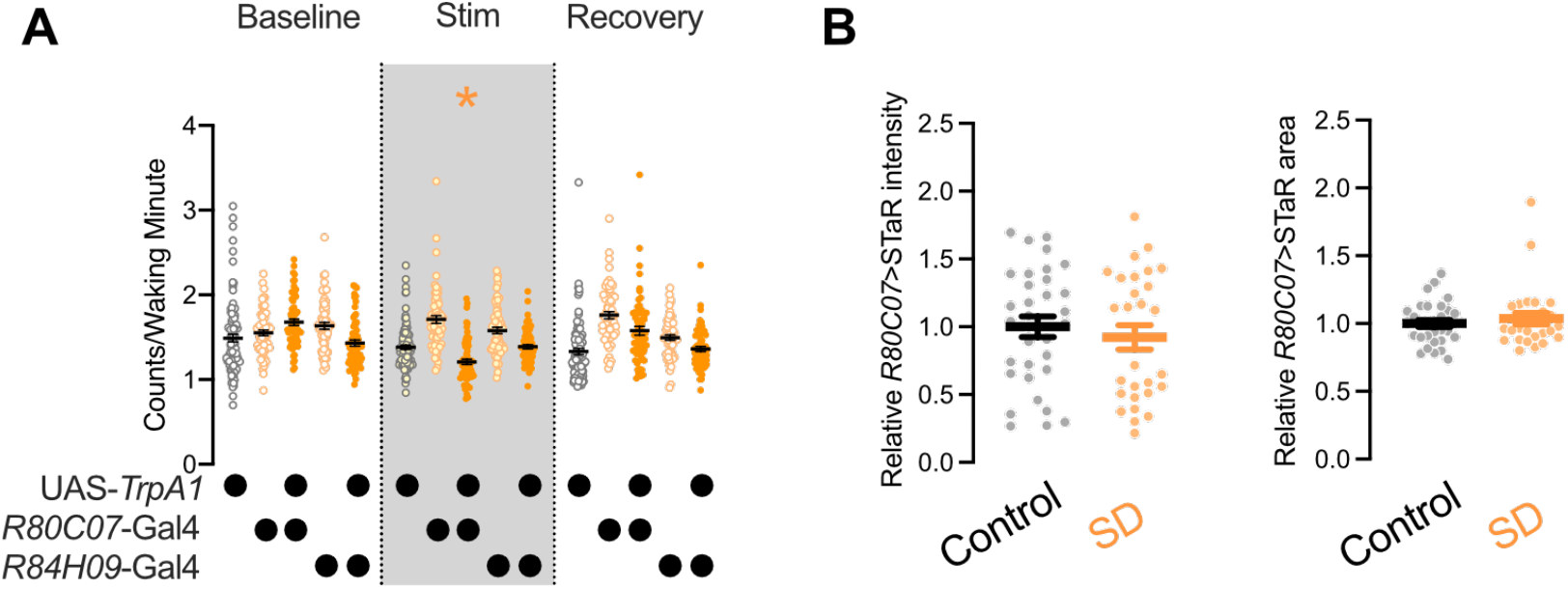
Waking activity during thermogenetic R3d activation; Related to Figure 3. **(A)** Waking activity (counts/waking minute) in experimental flies (orange) compared to genetic controls (Gal4/+ in light orange; UAS-*TrpA1*/+ in gray). Two-way repeated measures ANOVA finds a significant day-by-genotype interaction (F_(8,664)_=14.99, p<0.0001, n=59-93 flies/group). * signifies p<0.05 compared to Gal4/+ and UAS-TrpA1/+ controls by Tukey’s multiple comparison test). **(B)** Mean relative intensity (left) and cross-sectional area (right) of pre-synaptic BRP labelled by STaR expression in R3d neurons using *R80C07*-Gal4. Sleep deprivation had no significant effect on relative STaR intensity (Mann-Whitney test p=0.5412) or area (Two-tailed T-test p=0.4648, n=28-33 brains/group).

**Supplemental Figure 2.**
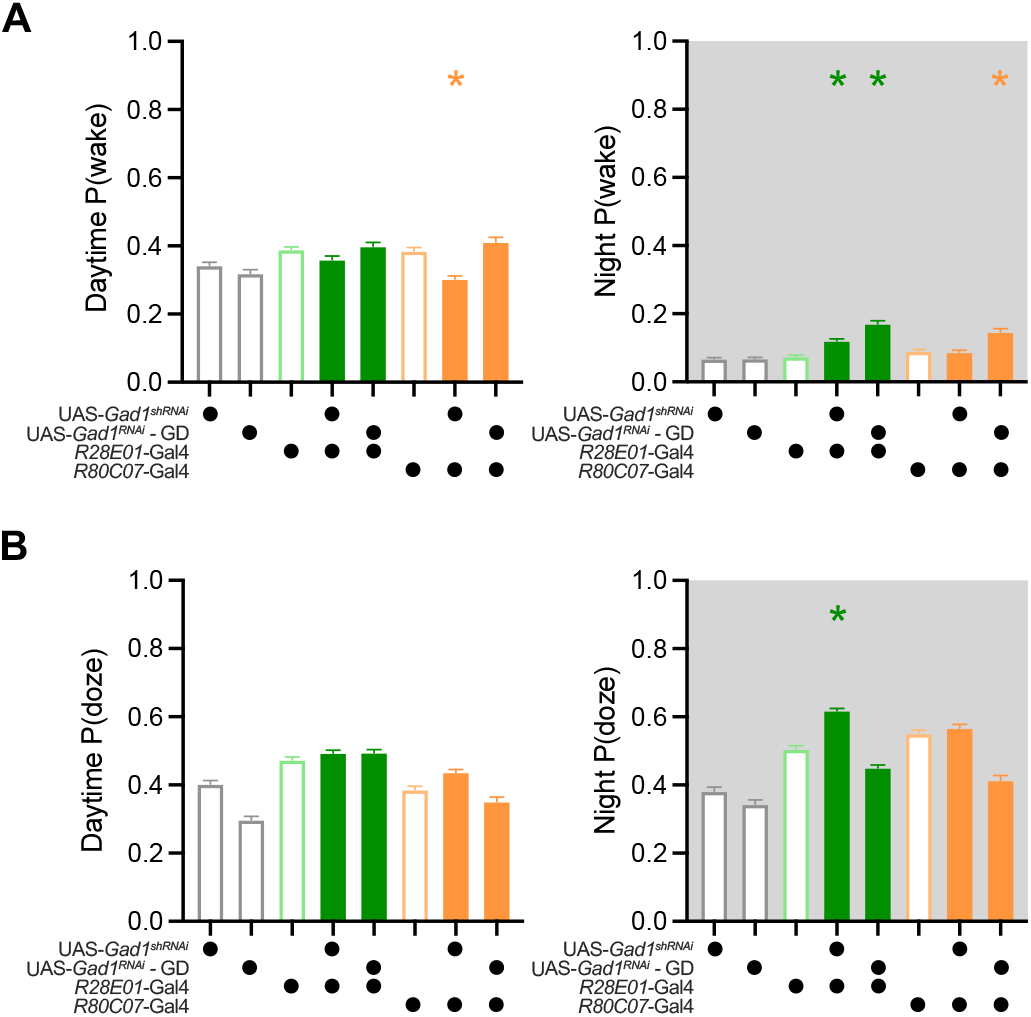
Effects of *gad1* knock-down in R3m and R3d neurons on sleep-wake state transitions; Related to Figure 4. **(A)** Minute-by-minute probability of waking from sleep (P(wake)) during the day (left panel) and night (right panel) for genotypes from **Figure 4C-F**. Daytime P(wake) is significantly reduced in *R80C07*-Gal4>UAS-*gad1*^shRNA^ flies; P(wake) is increased during the night when *R28E01-*Gal4 drives either RNAi construct and in *R80C07*-Gal4>UAS-*gad1*^RNAi^-GD flies. One-way ANOVA finds a significant genotype effect during the day (F_(7,1224)_=13.56, p<0.0001, n=92-192 flies/group) and night (F_(7,1225)_=37.12, p<0.0001, n=93-192 flies/group). **(B)** Probability that a waking fly will fall asleep (P(doze)) during the day (left) and night (right) for genotypes from **Figure 4C-F**. One-way ANOVA finds a significant effect for genotype during the day (F_(7, 1222)=_48.54, p<0.0001, n=92-192 flies/group) and night (F_(7, 1220)_=81.72, p<0.0001, n=92-192 flies/group).

**Supplemental Figure 3.**
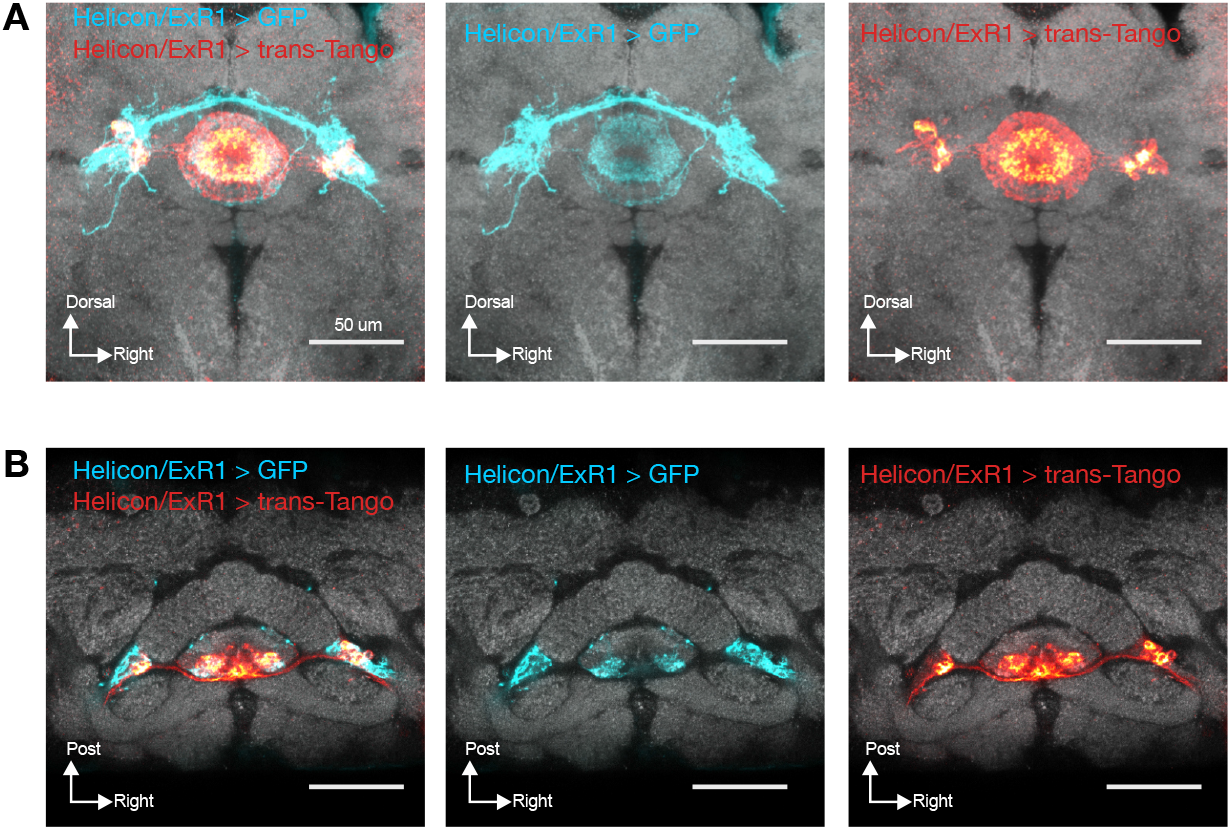
trans-Tango labeling of Helicon/ExR1 neuron post-synaptic targets; Related to Figure 5. **(A-B)** Labelling of Helicon/ExR1 neurons using a highly specific, intersectional split-Gal4 driver (Blue) along with trans-Tango tracing of post-synaptic targets (Red). Frontal view shown in **(A)**; dorsal view depicted in **(B)**

## STAR Methods

### RESOURCE AVAILABILITY

#### Lead Contact

Further information and requests for resources and reagents should be directed to and will be fulfilled by the Lead Contact, Jeffrey Donlea.

#### Materials Availability

This study did not generate new unique reagents.

#### Data and Code Availability

The published article includes all datasets generated during this study. This study did not generate any novel code.

### EXPERIMENTAL MODEL AND SUBJECT DETAILS

#### Fly Stocks and Maintenance

Fly stocks were reared on standard cornmeal media (per 1L H_2_O: 12g agar, 29g Red Star yeast, 71g cornmeal, 92g molasses, 16mL methyl paraben 10% in EtOH, 10mL propionic acid 50% in H_2_0) at 25°C with 60% relative humidity and entrained to a daily 12hr light, 12hr dark schedule. *Canton-S* were provided by Dr. Gero Miesenböck (University of Oxford), UAS-*TrpA1* (Hamada et al., 2008) were from Dr. Paul Shaw (Washington University in St. Louis). The following genetic driver lines were created for the Janelia Research Campus stock collection (Dionne et al., 2018; Jenett et al., 2012; Pfeiffer et al., 2010) and ordered from the Bloomington *Drosophila* Stock Center: *R12B01*-Gal4, *R12G08*-Gal4, *R28D01*-Gal4, *R28E01*-Gal4, *R31A12*-Gal4, *R34D03*-Gal4, *R37E10*-Gal4, *R47D08*-Gal4, *R58H05*-Gal4, *R59B10*-Gal4, *R78B06*-Gal4, *R80C07*-Gal4, *R84H09*-Gal4, *R92A09*-Gal4, *R24B11*-LexA, *R28E01*-LexA, *R48H04*-LexA, *R24B11*-p65.AD, and *R78A01*-Gal4.DBD. The following are Gal4 lines generated for the Vienna Tiles library (Tirian and Dickson, 2017) and generously shared by the Janelia Research Campus: *VT002226*-Gal4, *VT004309*-Gal4, *VT016270*-Gal4, *VT020036*-Gal4, *VT020036*-Gal4, *VT020613*-Gal4, *VT026873*-Gal4, *VT029750*-Gal4, *VT038873*-Gal4, *VT042805*-Gal4, *VT057232*-Gal4, *VT063740*-Gal4, and *VT063949*-Gal4. UAS-*Gad1*^RNAi^ stocks were acquired from the Vienna *Drosophila* Resource Center (VDRC; stocks 32344GD & 330039) (Dietzl et al., 2007), and the STaR effector line (w^-^; 20xUAS-RSR.PEST, 79C23S-RSRT-STOP-RSRT-smGFP_V5-2A-LexA/cyo; (Peng et al., 2018)) was shared by Dr. Orkun Akin (UCLA). nSyb-GRASP stocks (w*; P{w[+mC]=lexAop-nSyb-spGFP1-10}2, P{w[+mC]=UAS-CD4-spGFP11}2; MKRS/TM6B and w[*]; P{w[+mC]=UAS-nSyb-spGFP1-10}2, P{w[+mC]=lexAop-CD4-spGFP11}2/CyO; (Macpherson et al., 2015)), UAS-CD8::GFP (Pfeiffer et al., 2010), and LexAOP-mCD4::RFP (Pfeiffer et al., 2010) were provided by BDSC.

### METHOD DETAILS

#### Sleep

Sleep was measured as previously described (Shaw et al., 2002). Briefly, 3-7 day old female flies were individually loaded into 65mm-long glass tubes and inserted into *Drosophila* activity monitors (Trikinetics Inc; Waltham MA, USA). Periods of inactivity lasting at least 5 minutes were classified as sleep. Sleep deprivation occurred mechanically via the Sleep Nullifying Apparatus (SNAP) (Shaw et al., 2002). Trikinetics activity records were analyzed for sleep using Visual Basic scripts (Shaw et al., 2002) in Microsoft Excel or the Sleep and Circadian Analysis MATLAB Program (SCAMP) scripts (Donelson et al., 2012; Wiggin et al., 2020) in Matlab (Mathworks; Natick MA, USA). Multibeam monitoring experiments used MB5 monitors (Trikinetics Inc; Waltham MA, USA) to record fly movements.

#### Arousability

Arousability was tested by attaching Trikinetics activity monitors to microplate adapters on vortexers (VWR). Vibration force intensities were measured using Vibration 3.83 (Diffraction Limited Design; Southington CT, USA). Arousal tests used a 0.5g stimulation of 10 second duration.

#### Antennal Injury

Female flies were loaded into activity monitors at ∼4-7 days after eclosion, then permitted 1-2 days of baseline sleep. Antennae were bilaterally transected using fine forceps under CO_2_ anesthesia, then flies were returned to their tubes in activity monitors for recovery. Uninjured control siblings received matching CO_2_ concentration and exposure time.

#### Immunohistochemistry and confocal microscopy

*Drosophila* brains were dissected in PBS (1.86 mM NaH_2_PO_4_, 8.41 mM Na_2_HPO_4_, 175 mM NaCl; Sigma-Aldrich) and fixed in 4% (w/v) paraformaldehyde (Electron Microscopy; Hatfield PA, USA) in PBS for 30-45 minutes on ice. For GFP and RFP immunostaining, brains were incubated in primary antibody (1:1000 chicken anti-GFP, Molecular Probes; 1:1000 rabbit anti-DsRed, Takara Bio) overnight followed by secondary antibody (1:1000 anti-chicken conjugated to Alexa488, Molecular Probes; 1:1000 anti-rabbit conjugated to Alexa546, Molecular Probes) for roughly 24 hours. Immunostaining for V5 used a 48-hour incubation period in 1:400 mouse anti-V5 conjugated with DyLight550. For GRASP experiments, brains were incubated in primary antibodies (1:50 mouse anti-GFP, Sigma-Aldrich; 1:20 rat anti-N-cad, DSHB) for about 48 hours, followed by incubation in secondary antibodies (1:1000 anti-mouse conjugated to Alexa488, Molecular Probes; 1:1000 anti-rat conjugated to Alexa647, Jackson Immunoresearch Labs) for 48 hours. Brains stained for GABA were incubated in primary antibody (1:500 rabbit anti-GABA, Sigma-Aldrich) overnight, then overnight in secondary (1:1000 anti-rabbit conjugated with Alexa546, Molecular Probes).

Trans-Tango flies were reared at 18°C for 15-21 days post-eclosion, at which point they were dissected and fixed as described above, incubated in primary antibodies (as described above for GFP, RFP, and N-cad) overnight, followed by incubation in secondary antibodies (as described above for GFP, RFP, and N-cad) for 24 hours. All brains were mounted in Vectashield (Vector Labs; Burlingame CA, USA) and imaged on a Zeiss LSM880. All image processing was completed using Fiji (Schindelin et al., 2012).

Staining and mounting for genetic driver expression patterns in frontal and dorsal mount images (Figures 2A, 3A, 5) were conducted as previously described (Omoto et al., 2017, 2018).

### QUANTIFICATION AND STATISTICAL ANALYSIS

#### Statistical Analysis

Data were analyzed in Prism 9 (GraphPad; San Diego CA, USA). Group means were compared using two-tailed T-tests or one- or two-way ANOVAs, with repeated measures where appropriate, followed by planned pairwise comparisons with Holm-Sidak multiple comparisons tests. Sample sizes for each experiment are depicted in each figure panel or in the appropriate figure legend. All group averages shown in data panels depict mean ± SEM.

## Data Availability

All data generated during this study are included in the manuscript and supporting files.

## Notes

### Competing Interest Statement

The authors have declared no competing interest.

